# AFIDs-Validator: An Open-Access AI-Guided Platform for Learning Anatomical Landmark Placement

**DOI:** 10.64898/2026.08.20.746086

**Authors:** Alaa Taha, Dhananjhay Bansal, Jason Kai, Tristan Kuehn, Olivia W. Stanley, Patrick Park, Arun Thurairajah, Mackenzie Snyder, Greydon Gilmore, Mohamad Abbass, Borna Mahmoudian, Violet M. Liu, Jaime Thrower, Ali R. Khan, Jonathan C. Lau

**Affiliations:** Stanford University School of Medicine, Stanford, CA, United States of America; Imaging Research Laboratories, Robarts Research Institute, Western University, London, Canada; Child Mind Institute, New York, NY, USA; Department of Clinical Neurological Sciences, Division of Neurosurgery, Western University, London, Canada; Graduate Program in Neuroscience, Western University, London, Canada; Centre for Functional and Metabolic Mapping, Robarts Research Institute, The University of Western Ontario, London, Canada

**Keywords:** neuroimaging education, anatomical fiducials, spatial normalization, quality control, AI tutoring, large language models, brain atlas, MRI training, open science

## Abstract

Accurate localization of anatomical landmarks is a foundational skill in anatomy and imaging that is often taught informally through expert mentorship, requiring access to data and desktop software. There is no openly accessible, interactive resource that teaches neuroanatomy with quantitative feedback. We present the AFIDs-Validator (validator.afids.io), an open-access, browser-based platform that pairs guided instruction with quantitative assessment. The platform combines (1) a learning mode in which a language-model neuroanatomy tutor operates inside an MRI viewer, giving anatomy-first instruction that responds to the learner’s current image slice, orientation, and cursor position; and (2) a validation engine that accepts a learner’s landmark file and returns per-landmark Euclidean error against expert-annotated references spanning 21 brain templates. To make the feedback interpretable, we analyzed 15,000 landmark annotations across 132 human subjects and found that landmark difficulty varies fourfold (median error ranged from 0.37 mm at the anterior commissure to 1.50 mm at the temporal horns) with heavy-tailed distributions at every landmark. These distributions are compiled into per-landmark reliability priors, so learners are scored against the empirical spread of trained raters rather than an arbitrary threshold, and difficult landmarks are not mistaken for poor performance. The AFIDs-Validator requires no installation, licensed software, or local data, and all code, reference data, and tutor design are openly released.

## 1. Introduction

Reproducible neuroimaging depends not only on robust computational pipelines, but also on the ability of researchers and trainees to identify anatomy consistently. The past decade has produced a remarkable ecosystem of open-source infrastructure, including preprocessing pipelines such as fMRIPrep (Esteban et al., 2019) and FSL (Jenkinson et al., 2012), automated image-quality assessment through MRIQC (Esteban et al., 2017), data-sharing platforms such as OpenNeuro (Markiewicz et al., 2021), standardized template libraries through TemplateFlow (Ciric et al., 2022), and browser-native viewers such as NiiVue (Taylor & Rorden, 2023). These tools have lowered the barriers to reproducible, large-scale analysis. Yet interpreting their outputs still requires locating the same anatomical feature reliably across different brains of variable image quality, and in different coordinate spaces.

Locating a specific feature on a cortical sulcus, within the cerebral ventricles, or the exact crossing of a commissure on a grayscale volume is difficult and error prone, and the skill is most often taught through apprenticeship. A trainee typically learns through an expert, placing landmarks, and being corrected in real time. This model produces excellent raters but does not scale and may be unavailable to many trainees in lower resourced settings. Atlas reading and didactic lectures convey where structures are in the abstract but not how to find them on a specific noisy image, and they provide no direct feedback for the learner.

The Anatomical Fiducials (AFIDs) protocol was developed as a framework for landmark-based correspondence (Lau et al., 2019). It defines precisely specified landmarks distributed throughout the brain (from commissural midline structures to ventricular and sulcal features) with explicit operational definitions that minimize placement ambiguity. Original validation studies demonstrated mean inter-rater Euclidean errors of 1 to 2 mm and intraclass correlation coefficients (ICC) exceeding 0.9 for most landmarks after minimal training (Lau et al., 2019). Subsequent multi-cohort validation (Taha et al., 2023) established reliability benchmarks across 132 subjects, 30 rater sessions, and four imaging datasets, with rater experience spanning trainees with no prior imaging background to neurosurgical trainees. The AFIDs protocol is maintained as an open GitHub organization (github.com/afids) consisting of a written specification, a curated multi-rater dataset, and automated localization tools.

Despite the maturation of this protocol over the years, there is no interactive resource for learning landmark placement, and no browser-native way to check a placement against a reference. Existing workflows depend on desktop applications, including 3D Slicer (Fedorov et al., 2012), which has many capabilities and is extensible, but where installation overhead and version drift raise the barrier to both adoption and reproducibility. The gap is in the user-facing infrastructure that would make the AFIDs ecosystem, and the skill it encodes, broadly teachable. In this work, we present the AFIDs-Validator (validator.afids.io), which provides **(1) an AI-guided learning mode** that teaches landmark placement interactively with anatomical explanation, quantitative feedback, and no expert supervision, and **(2) an instant-feedback validation engine** for any AFIDs file against reference templates. This paper describes the platform, the pedagogical design of the AI tutor, the empirical basis for its feedback, and the accessibility choices that let a trainee use it end-to-end.

## 2. Methods

### 2.1 System overview and architecture

The AFIDs-Validator (see Figure 1) is a browser-based web application on a Flask 3.0 / Python 3.11 backend with a React 18 frontend. The validator is containerized with Docker Compose and deployed via Nginx, supporting both the hosted instance at: validator.afids.io and self-hosted institutional deployments. All source code is public at: github.com/afids/afids-validator under GPL-3.0.

**Figure 1.**
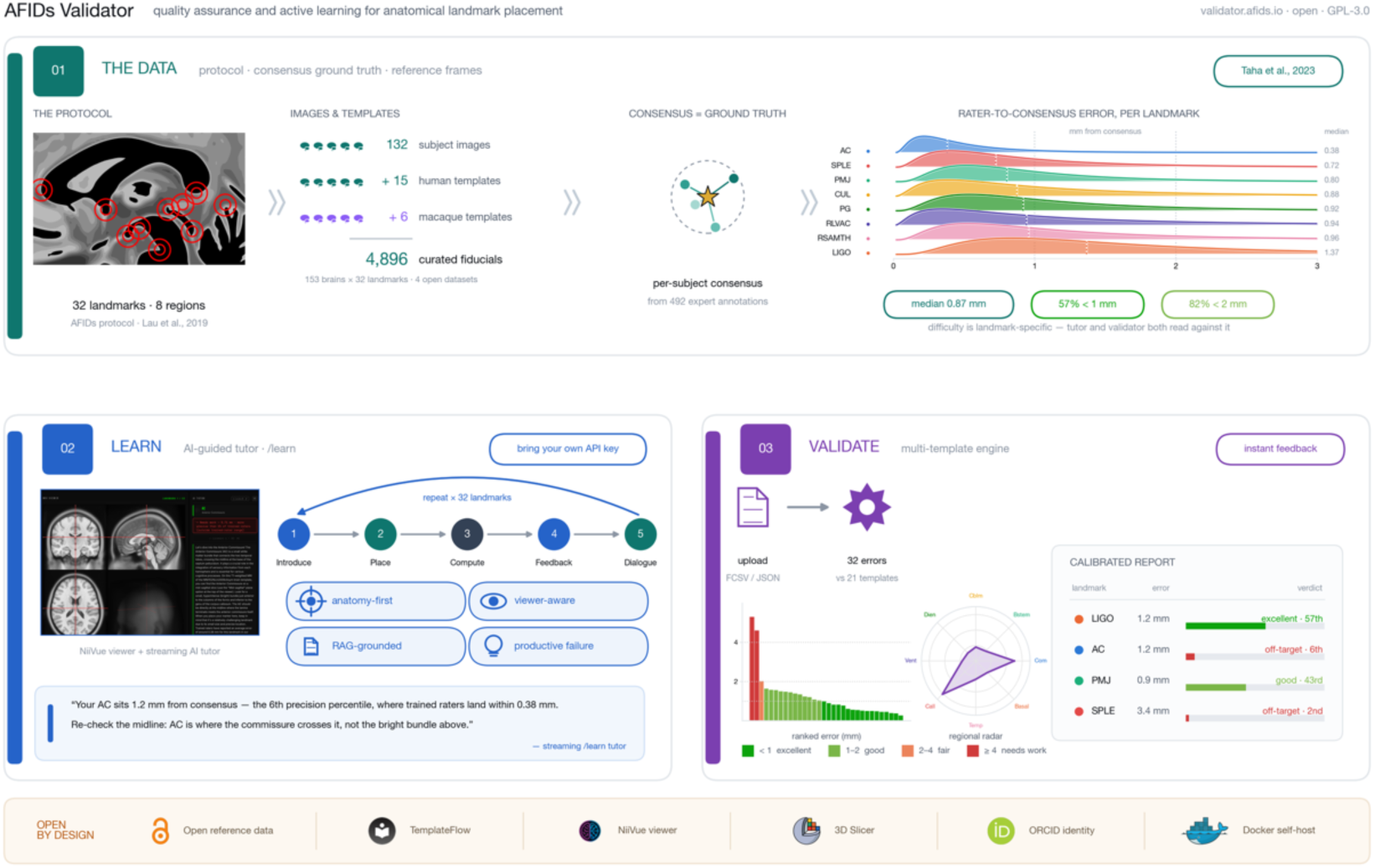
Architecture of the AFIDs-Validator. The platform comprises three modules. (1) Data foundation: trained-rater placements from the open AFIDs multi-rater release are distilled into per-landmark reference distributions of anatomical fiducial localization error (AFLE). (2) AI tutor: a browser-based guided-learning mode in which users place each of the 32 landmarks in an interactive viewer and receive protocol-grounded, reliability-calibrated feedback from a language model. (3) Validation engine: uploaded fiducial sets (FCSV) are scored against the reference, returning per-landmark error and session-level summary statistics. The platform is open by design: GPL-3.0 code, openly available reference data, and interoperability with TemplateFlow, NiiVue, 3D Slicer, ORCID, and Docker for self-hosting.

The complete guided-learning and validation workflows run in any modern browser on any operating system. ORCID OAuth (https://orcid.org) is optionally available for researcher identity, enabling longitudinal tracking of placement performance across sessions; authenticated users may opt in to contribute sessions to an institutional database (PostgreSQL via SQLAlchemy), and no placement data is retained without explicit consent.

The validator engine accepts two file formats. **FCSV** (Fiducial CSV) is the native export of 3D Slicer’s Markups module (Fedorov et al., 2012), which is the recommended placement tool in the AFIDs protocol. **JSON** is an alternative file produced by newer versions of 3D Slicer Markups module (Fedorov et al., 2012). Both are validated against the landmark schema, with automatic detection and internal conversion of coordinate conventions (i.e. RAS vs. LPS).

The learning and validation components of the AFIDs-Validator are deliberately continuous where a learner trains on the guided mode, then uploads an independent placement to the validation engine and receives the same quantitative feedback used in research quality control.

### 2.2 Guided learning mode

The guided learning mode (/learn) is the platform’s central pedagogical contribution (see Figures 2 and 3). This entails an interactive neuroanatomy training workflow for 32 anatomical landmarks (see Figure 4), in a browser. It embeds the NiiVue MRI viewer (Taylor & Rorden, 2023) loaded with the MNI152NLin2009cAsym T1w template from TemplateFlow (Ciric et al., 2022) beside a streaming LLM chat interface, a live coordinate readout, and per-landmark progress tracking.

**Figure 2.**
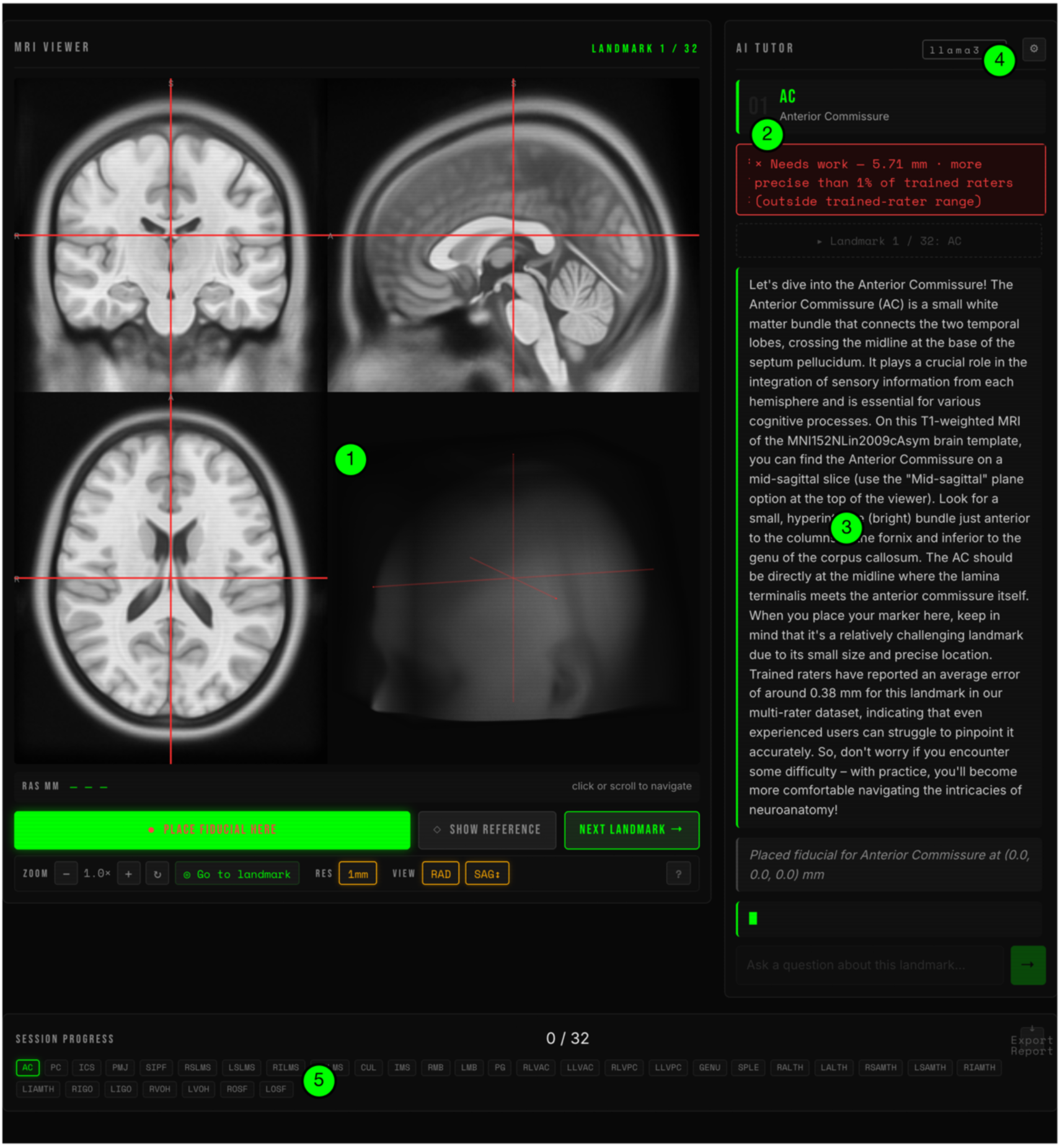
The guided-learning interface (live session). Annotated screenshot of the /learn mode running against a local model. (1) the browser-based NiiVue MRI viewer (MNI152NLin2009cAsym T1w), here showing a placed fiducial for the anterior commissure across the three orthogonal planes and a 3-D rendering; (2) the rater-calibrated result badge reporting Euclidean error and the learner’s percentile within the trained-rater distribution for this landmark; (3) the difficulty-aware, anatomy-first AI tutor, whose feedback cites the landmark’s real trained-rater difficulty in plain language; (4) the active model and bring-your-own-key control; (5) the 32-landmark progress tracker and session-report export.

**Figure 3.**
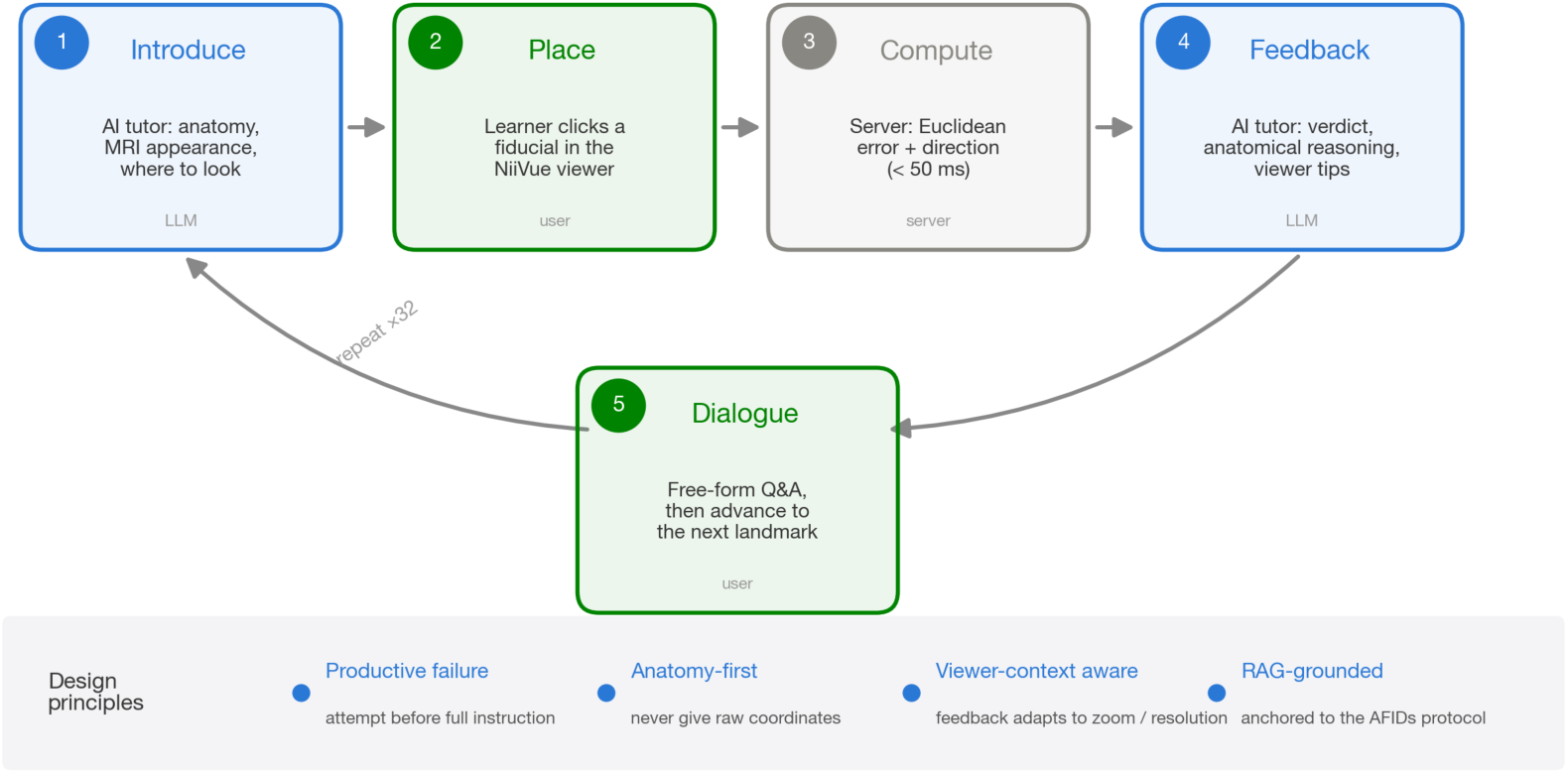
The guided-learning cycle and its design principles. Each landmark proceeds through five steps: (1) AI introduction; (2) learner placement in the viewer; (3) server-side error computation (<50 ms); (4) anatomy-first AI feedback; (5) free-form dialogue; the cycle repeats across all 32 landmarks. The principles governing the tutor are annotated below: productive failure (attempt before full instruction), anatomy-first (never give raw coordinates), viewer-context awareness (feedback adapts to zoom/resolution), and retrieval-augmented grounding in the AFIDs protocol.

**Figure 4.**
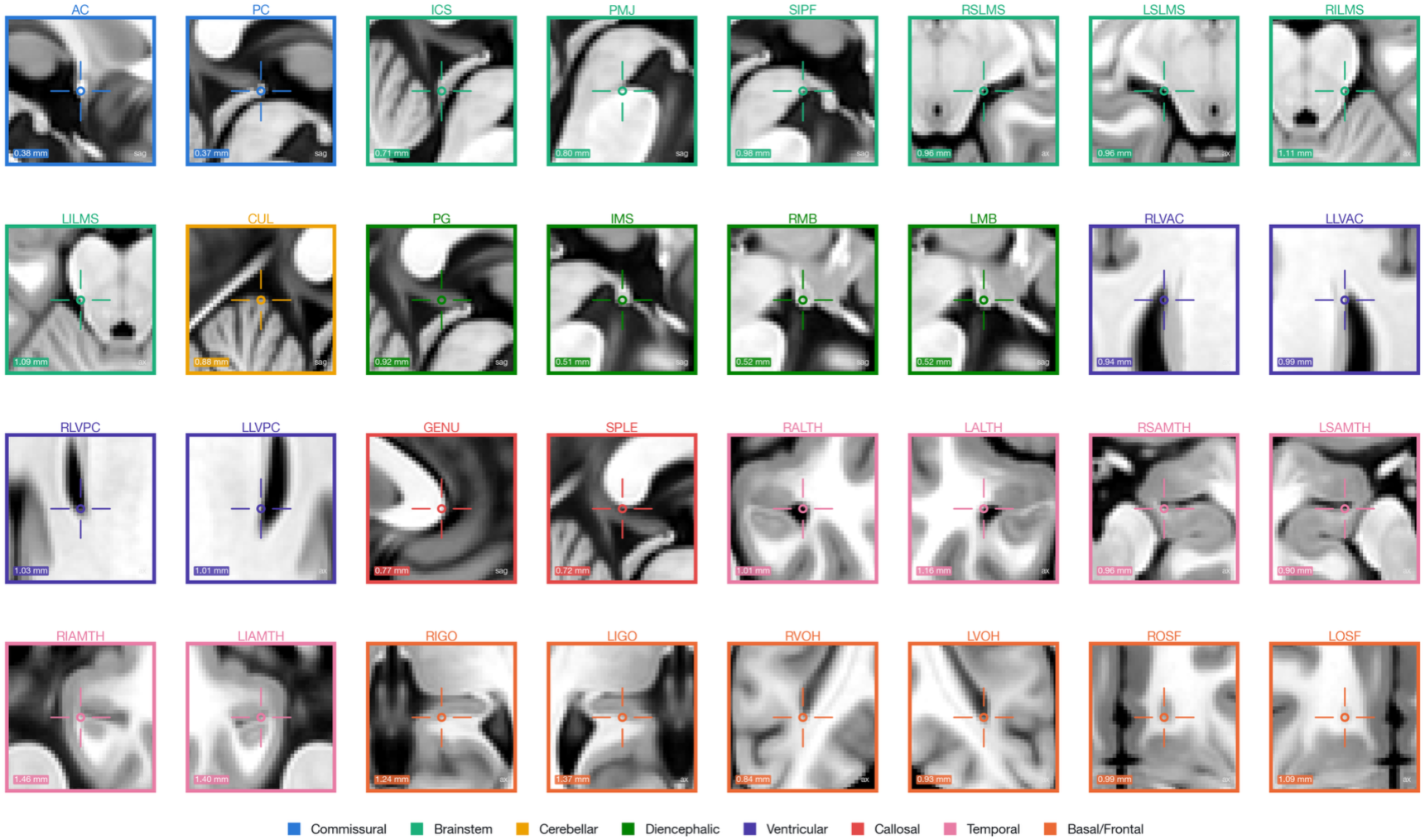
A guide to the 32 AFIDs landmarks on the MNI152NLin2009cAsym T1w template. Each panel is a 40 mm patch of the template centered on one landmark, with a crosshair marking the exact point, grouped and colored by neuroanatomical region. The chip in each panel gives that landmark’s median trained-rater localization error (AFLE) and the corner tag its viewing plane (sag = sagittal, used for midline landmarks; ax = axial, for bilateral landmarks).

Each landmark proceeds through five steps:

1. **Introduction** (LLM, streamed, less than 5 sentences): the structure’s anatomical identity and functional significance; its appearance on T1w contrast; the recommended imaging plane; and the single most common placement error.
2. **Placement** (learner): a click in NiiVue records a coordinate.
3. **Computation** (server, <50 ms): Euclidean distance to the template reference, directional offsets, and a quality rating.
4. **Feedback** (LLM, streamed, ≤6 sentences): a one-line verdict; anatomical reasoning about the likely error; directional guidance in anatomical language; and, when relevant, a specific viewer adjustment.
5. **Dialogue** (learner-initiated): free-form Q&A with maintained conversation history.

A learner completing the full set produces a downloadable session report containing a per-landmark table of placements, distances, and quality, together with the full tutoring transcript that serves both as a study artifact and as documentation of proficiency. Well-designed intelligent tutoring systems can approach the effectiveness of one-on-one human tutoring when they pair step-level feedback with sound instructional design (VanLehn, 2011); we accordingly design the tutor’s behavior around a small set of principles from the learning sciences:

### Contextual instruction

The learner encounters each landmark in situ, where instruction arrives at the moment of placement, in the same viewer, on the same image they will be evaluated against. Situating learning in the environment of use improves retention and transfer relative to decontextualized atlas study (Lave & Wenger, 1991; Koedinger & Corbett, 2006).

### Active generation before instruction

The mode gives a brief orientation but withholds full explanation until the learner has attempted a placement. This ordering reflects the “productive failure” framework (Kapur, 2008, 2016; Loibl et al., 2017): learners who struggle with a problem before receiving targeted instruction show superior long-term retention and transfer. A trainee who has tried to locate the posterior commissure integrates the subsequent explanation differently than one who merely read it.

### Anatomy grounded explanations

The tutor’s system prompt explicitly forbids giving target coordinates or numerical navigation unless explicitly asked (e.g., “give me the coordinates of the thalamus”). This defends against a common failure mode of LLMs used as anatomical assistants which train lookup rather than recognition and produces a skill that collapses on individual-subject data with variable anatomy.

### Viewer-context-aware scaffolding

The NiiVue viewer state at the moment of placement (e.g., zoom, image resolution, and contrast window) is captured and included in the feedback request, letting the tutor recommend concrete viewer changes: *“you placed this at low zoom in 2 mm resolution; switching to 1 mm via the resolution (“RES”) button and zooming in would reveal the fine structure here“*. This adaptation to the learner’s visual environment can provide the guidance necessary for improving accuracy.

### Scaffolded, protocol-grounded feedback

Rather than embedding all 32 landmark definitions in every prompt, the tutor is grounded by retrieval-augmented generation (RAG): each AFIDs landmark definition is embedded and stored, and for a given landmark or free-text question the most semantically relevant definitions are retrieved and injected into the model’s context. This is complemented by a separate rater-reliability signal: for each placement, the learner’s error is compared against the distribution of trained-rater localization error for that landmark, and a calibration statement (how the placement ranks relative to expert raters) is injected directly into the prompt so that feedback is calibrated to how difficult each landmark actually is.

The AI tutor’s language-model engine is configurable through environment variables (LLM_API_KEY, LLM_BASE_URL, LLM_MODEL). It supports commercial models (e.g., GPT-4o; Anthropic Claude via a compatible endpoint), locally hosted open-weight models (e.g., Llama 3, Mistral via Ollama), and low-cost hosted providers. To remove cost as a barrier, the platform defaults to Groq’s free Llama 3.3 70B endpoint, and users can enable the tutor with a free API key obtained in minutes (step-by-step instructions are in the project README at github.com/afids/afids-validator); the key is stored only in the user’s browser and is never logged server-side. Institutions without commercial API access can run a fully functional tutor on local GPU resources via Ollama. The pedagogical value derives from the structured prompt, RAG grounding, and feedback workflow rather than any single model’s capabilities.

NiiVue is embedded as a WebGL2 canvas. The MNI152NLin2009cAsym T1w volume is fetched from TemplateFlow’s public S3 bucket on first access and cached server-side, at 2 mm (∼1.7 MB) and 1 mm (∼9 MB, default) isotropic. Placement coordinates are returned via the NiiVue API to the Flask /learn/check endpoint. Viewer state (zoom, resolution, contrast window) is captured at placement and included in the feedback request. LLM communication uses the OpenAI-compatible Python SDK (≥1.0) with streaming over chunked HTTP; per-request overrides (api_key, base_url, model) supplied by the client take precedence over the server default and are neither logged nor persisted. Landmark context is assembled by retrieval-augmented generation over an embedded knowledge store of AFIDs definitions and protocol passages, with fallback to the curated landmark dictionary.

### 2.3 Validation engine

For each of the 32 landmarks the validator computes:

1. **Euclidean distance** from the user coordinate to the template reference (mm);
2. **Directional decomposition**: left/right, anterior/posterior, superior/inferior components (mm), suppressing components below 0.5 mm;
3. **Quality classification:** excellent (<1 mm), good (1–2 mm), fair (2–4 mm), needs work (≥4 mm).

Session-level statistics across all landmarks are reported: median and mean error, standard deviation, best- and worst-performing landmarks, and counts within 1 mm and 2 mm. The four-tier quality classification communicated to the user is anchored to the published AFIDs multi-rater reliability dataset (Taha et al., 2023). That dataset comprises 132 subjects across four cohorts, with 30 rater sessions spanning novice (0 months imaging experience) to expert (≥24 months), accumulating >300 rater-hours and >45,000 Euclidean-distance measurements. In that dataset, trained raters achieved mean inter-rater errors of 1.0 to 1.5 mm for most landmarks, with ICC > 0.9 for landmarks including AC, PC, and the mammillary bodies. The 2 mm boundary corresponds to roughly the 82nd percentile of trained-rater errors (see section 3.1), a realistic proficiency target. Errors exceeding 4 mm corresponded to anatomical confusion events (e.g., placing on the habenular commissure instead of PC, or the corpus-callosum body instead of the splenium) rather than imprecision. The thresholds therefore discriminate distinct performance phenotypes rather than carving an arbitrary continuum, which is what makes them meaningful as feedback to a learner.

Three complementary Plotly visualizations are generated per session (Figure 5): a 3D scatterplot of template and user landmark sets with error-colored connecting lines (localizing spatial error patterns, e.g., a global lateral shift indicating a convention mismatch); a ranked error histogram identifying outlier landmarks for targeted practice; and a regional radar chart revealing region-level biases that implicate specific anatomical confusions. Parsing handles FCSV (Markups v4.6+) and AFIDs JSON. Structural validation confirms all 32 labels present, numeric coordinates, and a valid CoordinateSystem header. Accuracy of placements is validated via Euclidean distance at the following thresholds calibrated to Taha et al. (2023): excellent <1 mm, good 1–2 mm, fair 2–4 mm, needs work ≥4 mm, and refined per landmark by the rater-reliability prior (§2.5).

**Figure 5.**
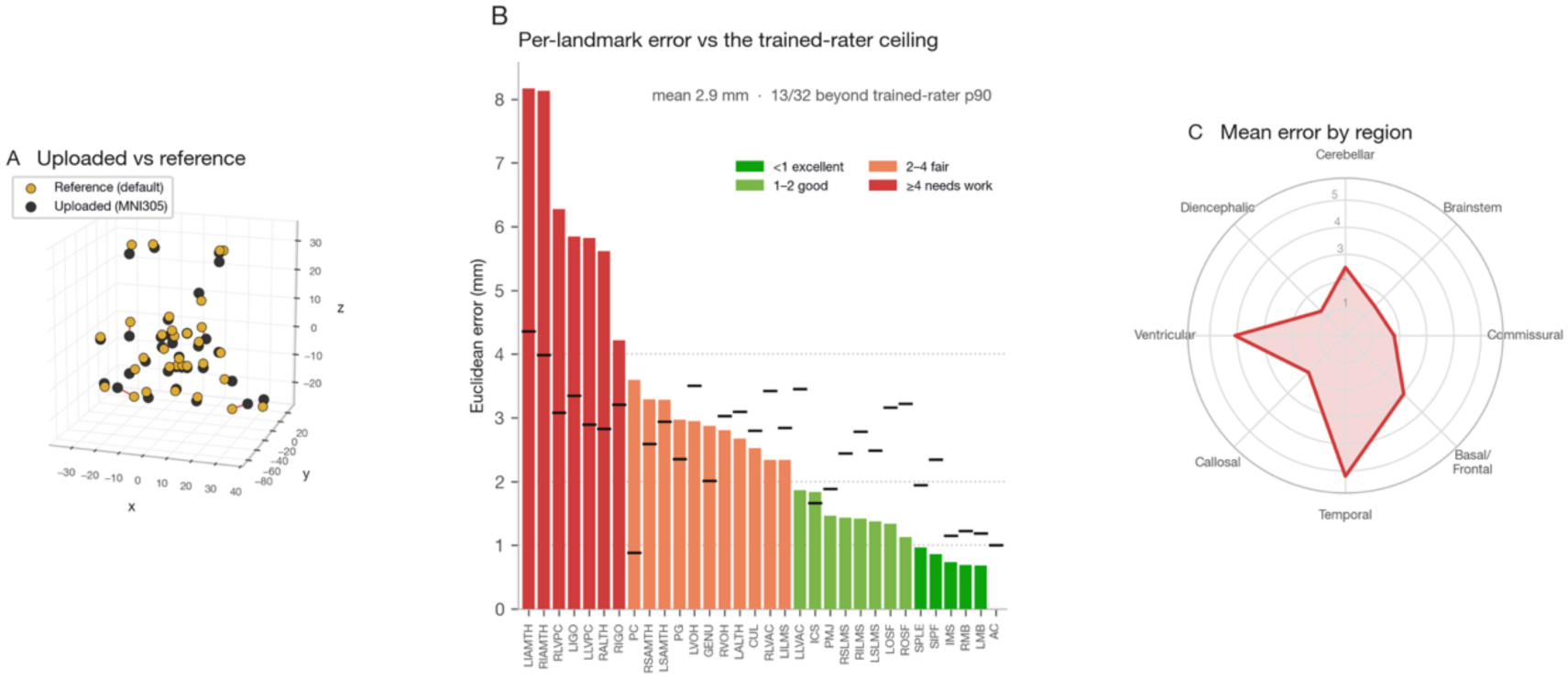
A worked quality-control catch on real reference data. The validation engine run on a genuine template-space mismatch: the MNI305 landmark set validated against the platform default (MNI152NLin2009cAsym). (A) 3D scatter of the default reference (gold) and the uploaded MNI305 landmarks (black), connected by lines colored by error magnitude. (B) Per-landmark Euclidean error, ranked and colored by quality tier (excellent <1 mm, good 1–2 mm, fair 2–4 mm, needs work ≥4 mm); black ticks mark each landmark’s trained-rater 90th percentile, the error exceeds it for 13 of 32 landmarks (mean 2.9 mm), most dramatically at the posterior commissure (3.6 mm, 4.1× its rater 90^th^ percentile). (C) Mean error per neuroanatomical region: the signature is localized to the temporal and ventricular landmarks while the diencephalic floor stays tight.

### 2.4 Reference template library

All reference FCSV files were generated by expert raters following the AFIDs protocol (https://afids.github.io/afids-protocol/) and version-controlled in the AFIDs GitHub organization. FCSV files encode coordinates in RAS convention (3D Slicer v4.6+); the validator reads the CoordinateSystem header and converts to a canonical internal representation. All 21 templates contain all 32 landmarks.

The reference library contains 21 fully annotated brain atlases (15 human and 6 macaque) each with all 32 AFIDs landmarks, yielding 480 annotated human and 192 macaque ground truth landmarks (672 in total). All reference FCSV files are version-controlled in the AFIDs GitHub organization and distributed with the validator.

All 21 reference templates (15 human and 6 macaque) are catalogued together in Table 1. The human set spans the major contemporary standards, from MNI305 (Collins et al., 1994) to the 20-µm BigBrain histological atlas (Amunts et al., 2013), with complete coverage of the MNI152 family that serves as the default output space of major pipelines, plus the Parkinson’s-optimized PD25 (Xiao et al., 2017). The six macaque templates are D99 (Reveley et al., 2017), INIA19 (Rohlfing et al., 2012), MacaqueMNI (Frey et al., 2011), NMTv1.3 (Seidlitz et al., 2018), NMTv2.0asym (Jung et al., 2021), and Yerkes19 (Donahue et al., 2016).

**Table 1.** Brain templates in the AFIDs-Validator reference library (15 human, 6 macaque; all with the full 32-landmark set)

| Template | Species | Primary use | Citation |
| --- | --- | --- | --- |
| MNI152NLin2009cAsym | Human | fMRIPrep default; contemporary standard | Fonov et al., 2011 |
| MNI152NLin2009cSym | Human | Symmetric variant; FreeSurfer normalization | Fonov et al., 2011 |
| MNI2009cAsym | Human | Near-duplicate alias of MNI152NLin2009cAsym | Fonov et al., 2011 |
| MNI152NLin2009bAsym/Sym | Human | Legacy MNI152 2009b variants | Fonov et al., 2011 |
| MNI152NLin6Asym/Sym | Human | FSL standard space; diffusion imaging | Fonov et al., 2009 |
| MNI152Lin | Human | SPM linear normalization | Fonov et al., 2011 |
| MNI305 | Human | Original MNI standard; clinical reference | Collins et al., 1994 |
| Colin27 | Human | High-resolution single-subject MNI template | Holmes et al., 1998 |
| BigBrain | Human | 20-μm isotropic histological atlas | Amunts et al., 2013 |
| fsaverage | Human | FreeSurfer average surface template | Fischl et al., 1999 |
| OASIS30ANTs | Human | Multi-subject aging template (ANTs registration) | Marcus et al., 2007 |
| PD25 | Human | Parkinson's disease cohort template | Xiao et al., 2017 |
| Agile12v2016 | Human | Ultra-high field MRI Population template | Lau et al., 2019 |
| D99 | Macaque | Histology-based rhesus stereotaxic atlas | Reveley et al., 2017 |
| INIA19 | Macaque | Primate parcellation & spatial normalization | Rohlfing et al., 2012 |
| MacaqueMNI | Macaque | Macaque MNI-style population template | Frey et al., 2011 |
| NMTv1.3 | Macaque | NIMH Macaque Template v1.3 | Seidlitz et al., 2018 |
| NMTv2.0asym | Macaque | NIMH Macaque Template v2.0 (asymmetric) | Jung et al., 2021 |
| Yerkes19 | Macaque | Surface-based macaque population template | Donahue et al., 2016 |

### 2.5 Rater-reliability prior

Per-landmark trained-rater reliability was computed from the AFIDs multi-rater release (Taha et al., 2023). For each subject with multiple rater placements, each rater’s Euclidean distance to the per-subject consensus (ground truth) was measured for every landmark and aggregated across all four released cohorts (AFIDs-HCP, AFIDs-OASIS, SNSX, and LHSCPD; 492 rater files, 132 subjects) to obtain, per landmark, the median, mean, and 10th/25th/50th/75th/90th percentiles of AFLE. At run time the tutor maps a learner’s per-landmark error to a percentile within this distribution and a four-level band (better than typical / within range / edge / outside), injected into the feedback prompt and returned by /learn/check; landmarks absent from the prior fall back to the global fixed thresholds.

### 2.6 Inter-template variability analysis

The analysis was restricted to the eight canonical MNI152/MNI305 templates (MNI152Lin; MNI152NLin2009bAsym/bSym/cAsym/cSym; MNI152NLin6Asym/6Sym; MNI305), so that variability is measured within a directly comparable family. AC-normalized coordinates were computed by subtracting each template’s AC coordinate from all 32 positions. For each of the 31 non-AC landmarks we computed the cross-template centroid, each template’s Euclidean distance from it (mean ± SD), and the maximum pairwise distance.

## 3. Results

The reference library answers two quantitative questions that ground the platform’s two major contributions. First, landmark localization difficulty is computed from 492 expert placements, which tells the guided-learning tutor what a good attempt looks like landmark by landmark.

Second, we present a worked quality-control example on the reference templates themselves. We deliberately foreground these over the descriptive template statistics because what matters for a learner is exactly how much difficulty each landmark carries and whether the engine flags a mistake a practitioner would make.

### 3.1 A difficulty spectrum for landmark placement

Some landmarks are harder to identify than others, and the guided-learning mode is only as good as its model of that difficulty. We quantified this property more directly using the released multi-rater dataset: for every subject with multiple raters, each rater’s Euclidean distance to the per-subject consensus, the anatomical fiducial localization error (AFLE), was measured for all 32 landmarks and aggregated across all four cohorts (492 rater files, 132 subjects; Figure 6A). This difficulty analysis draws on the human multi-rater dataset; the six macaque templates serve as validation references and were not included in the rater-reliability analysis.

**Figure 6.**
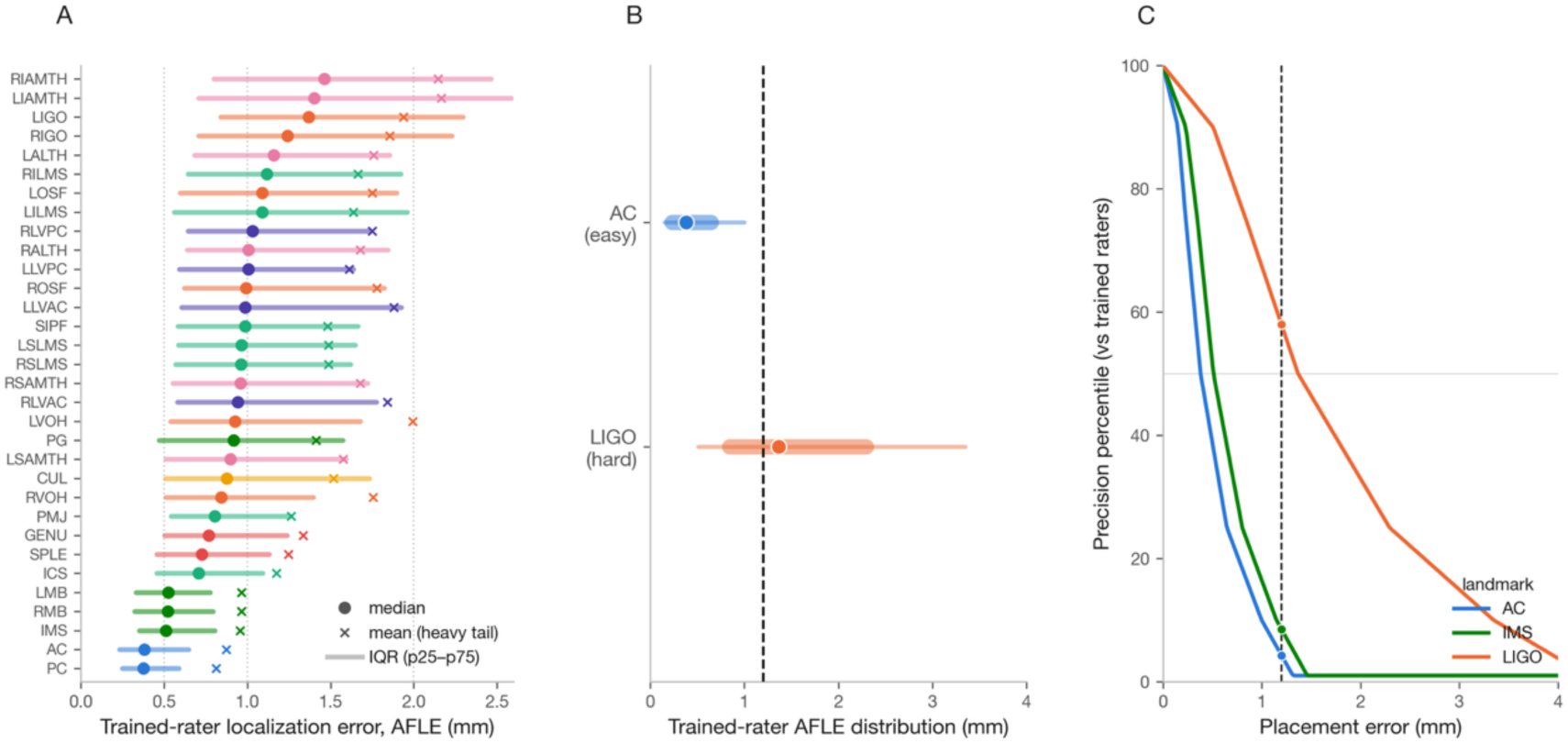
A difficulty benchmark from 492 expert placements. Computed from 492 trained-rater annotations across all 132 subjects of the AFIDs multi-rater release (Taha et al., 2023). (A) Per-landmark trained-rater localization error (AFLE), each landmark shown as its median (dot), interquartile range (bar), and mean (cross), ranked from most reliable (PC, AC; median ∼0.37 mm) to least (temporal-horn landmarks and indusium griseum origins; median up to ∼1.5 mm) which is a fourfold spread; the mean sitting right of the median at every landmark shows the heavy tail. (B) The same 1.2 mm placement, two verdicts: read against each landmark’s trained-rater distribution, 1.2 mm falls at roughly the 4th precision percentile for the anterior commissure (outside the expert range) but near the median (∼58th precision percentile) for the indusium griseum origin (a solidly expert placement). (C) The mm-to-percentile calibration the platform applies, shown for an easy (AC), typical (IMS), and hard (LIGO) landmark; the same error maps to sharply different precision percentiles.

Median AFLE varies approximately fourfold across the protocol (Figure 6A). The lowest median AFLEs occur at the commissures (PC, 0.37 mm; AC, 0.38 mm) and along the diencephalic floor, including the mammillary bodies and intermammillary sulcus (∼0.5 mm). In contrast, the highest median AFLEs occur at the temporal-horn landmarks (RIAMTH, 1.46 mm; LIAMTH, 1.40 mm) and the indusium griseum origins (LIGO, 1.37 mm; RIGO, 1.24 mm), which lie along thin, CSF-adjacent structures with less clearly defined boundaries. The higher AFLEs at these landmarks likely reflect both greater localization difficulty and greater underlying anatomical variability, rather than localization difficulty alone. Across all landmarks, the global median AFLE is 0.87 mm, with 57% of trained-rater placements falling within 1 mm and 82% within 2 mm.

Importantly, the error distribution is heavy-tailed for every landmark. Even for the commissures, which have a median AFLE of approximately 0.37 mm, the mean error is roughly 0.8 mm (about 2.2 times the median); across landmarks, the mean ranges from 1.4 to 2.3 times the median (Figure 6A, mean markers). Because AFLE is each rater’s distance to the per-subject consensus, measured within that subject’s own image, it reflects how reliably raters localize a landmark which is a mix of the landmark’s intrinsic ambiguity and occasional misidentification. The right tail therefore flags placements far from consensus, whether from a genuinely ambiguous landmark or an outright misidentification. Accordingly, the tutor treats a placement beyond a landmark-specific high percentile of the trained-rater distribution as a likely misidentification rather than mere imprecision, applying a per-landmark tolerance rather than a single global threshold.

### 3.2 Calibrated feedback relative to trained raters

Our platform evaluates performance using the empirical error distribution for each landmark. As shown in Figure 6B, the same 1.2 mm placement can have different meanings depending on the target. At the anterior commissure, this places the learner beyond essentially the entire trained-rater range for the anterior commissure (∼4th precision percentile, outside the expert range). At the indusium griseum origin, however, the same 1.2 mm falls near the median, corresponding to approximately the 58th precision percentile, potentially reflecting expert-level placement.

This landmark-specific reliability prior allows the AI tutor to grade each placement relative to the trained raters who defined that landmark. After each attempt, the validator reports the learner’s percentile and assigns one of four bands relative to that landmark’s trained-rater error distribution: 1) better than the typical trained rater, 2) within the trained-rater range, 3) at the edge of that range, or 4) outside it. Figure 6C shows the underlying calibration between millimeter error and percentile. This approach converts an otherwise abstract distance into a meaningful, landmark-specific benchmark that is also included in the session report. Because the prior is recalculated from the released placement data, it can be refined as the dataset grows. This underscores a central design principle of the AFIDs-Validator: an absolute placement error is only interpretable relative to the specific landmark it targets — precisely what the platform’s per-landmark reliability priors provide.

### 3.3 Two kinds of difficulty across templates

Rater difficulty (how hard a landmark is to *localize* on one image) is related to but distinct from inter-template variability (how much it *moves* across reference templates). Across the 31 non-AC landmarks the two correlates at **r = 0.66** (Figure 7A): hard-to-localize landmarks tend also to vary across templates, yet neither predicts the other completely — the ventral occipital horns, for instance, vary substantially across templates but are placed reliably by raters. A learner benefits from knowing which kind of difficulty a landmark carries. The reliability itself is balanced bilaterally: the mean left–right difference in median AFLE is only **+0.03 mm** across the 11 homologous pairs.

**Figure 7.**
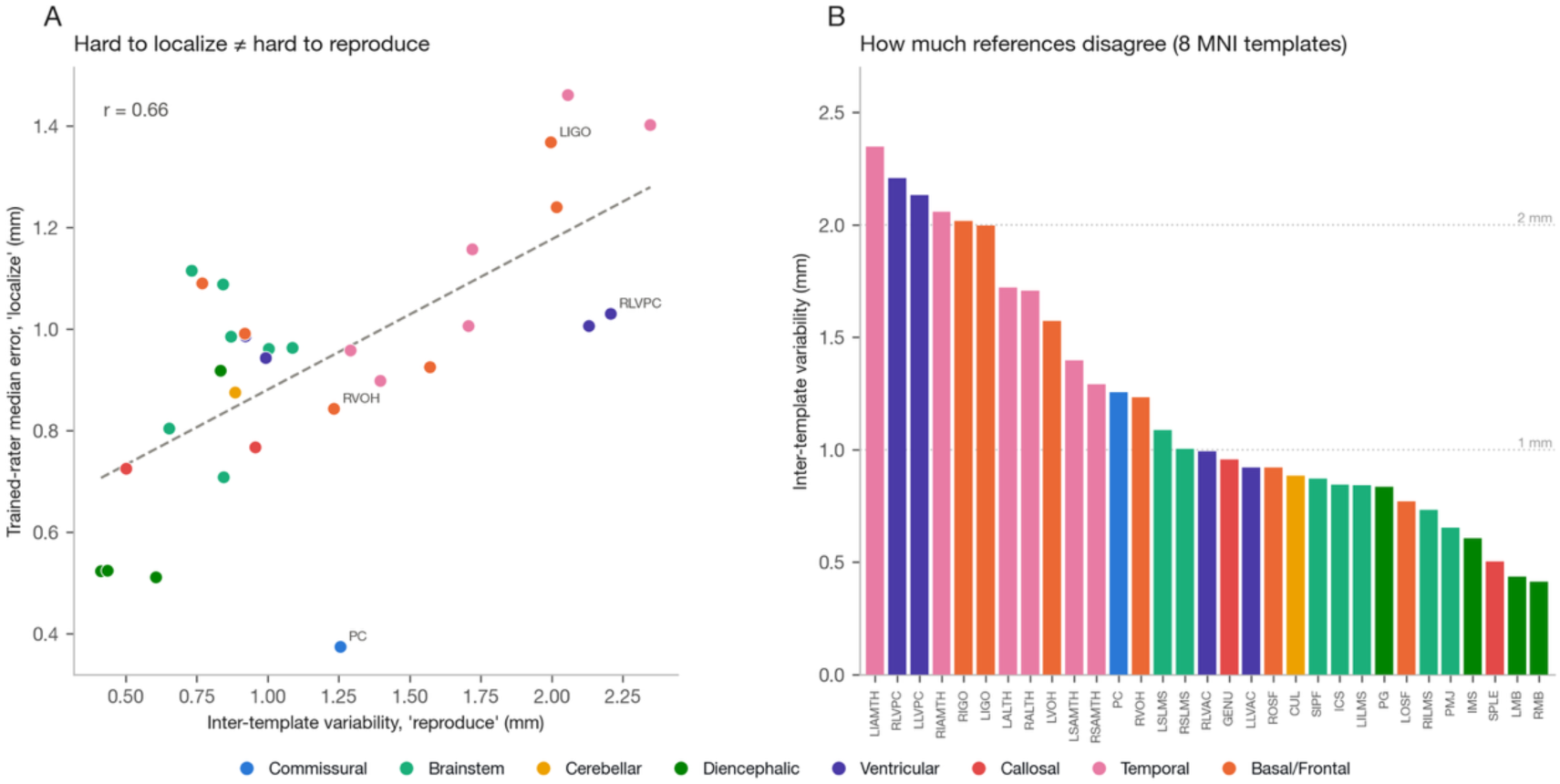
Two distinct kinds of difficulty. (A) Per-landmark trained-rater median error (“localize”) vs. inter-template variability (“reproduce”) for the 31 non-AC landmarks (Pearson’s r = 0.66): related but distinct, with labelled points that diverge from the trend (e.g., the ventral occipital horn RVOH varies across templates yet is placed reliably by raters). (B) Per-landmark AC-normalized inter-template variability across the eight canonical MNI152/MNI305 templates (mean distance to the cross-template centroid), ranked from the temporal-horn/ventricular landmarks (LIAMTH 2.35 mm) down to the mammillary bodies (RMB 0.41 mm), colored by region; dotted lines mark the validator’s 1 and 2 mm thresholds.

Expressing each template’s landmarks relative to its own AC and restricting to the eight canonical MNI152/MNI305 templates, AC-normalized variability ranges from 0.41 mm (right mammillary body) to 2.35 mm (left temporal horn), mean 1.20 mm, ranked by region in Figure 7B and Table 2. AC–PC distance spans 27.8–31.0 mm: the modern nonlinear MNI152 variants cluster within ∼28.0 ± 0.2 mm, while the linear MNI152Lin (30.3 mm) and original MNI305 (31.0 mm) diverge by 2–3 mm. The practical consequence is that references are not freely interchangeable for every landmark, which motivates the template-specific references.

**Table 2.** Inter-template variability by neuroanatomical region (AC-normalized, 8 MNI templates)

| Region | N landmarks | Mean variability (mm) | Max pairwise (mm) |
| --- | --- | --- | --- |
| Diencephalic | 4 | 0.57 | 3.67 |
| Callosal | 2 | 0.73 | 3.60 |
| Brainstem | 7 | 0.86 | 3.63 |
| Cerebellar | 1 | 0.89 | 3.65 |
| Commissural | 1 | 1.26 | 3.89 |
| Basal/Frontal | 6 | 1.42 | 8.29 |
| Ventricular | 4 | 1.56 | 6.61 |
| Temporal | 6 | 1.75 | 9.65 |

### 3.4 A quality-control example

We reproduced a rather well-known misalignment profile across the MNI template family. More specifically, landmarks defined in one MNI space are checked against a reference in another, by validating the MNI305 landmark set against the platform default (MNI152NLin2009cAsym), aligned only at the anterior commissure so that the residual is the geometry a proper registration would still have to recover (Figure 5). The engine reports a mean error 2.9 mm, with 13 of 32 landmarks beyond their own trained rater 90th percentile and the error concentrated in the temporal (5.2 mm) and ventricular (4.1 mm) regions while the diencephalic floor stays tightly aligned (1.3 mm). The posterior commissure alone lands at 3.6 mm (4.1 times its trained rater 90th percentile). Critically, the engine grades severity: repeating the exercise with the milder MNI152Lin mismatch yields a mean of only 1.6 mm with 23 of 32 landmarks still within 2 mm, correctly reading as a subtle rather than gross discrepancy. This region-resolved, per-landmark verdict is precisely what a global image-similarity score or a visual pass may miss, and it is the same output a learner receives on their own upload.

## 4. Discussion

### 4.1 Accessibility, equity, and reproducibility

The AFIDs-Validator runs in any modern browser with no local installation. Learners can run the tutor on a shared default, on a free or low-cost hosted model, or on a locally hosted open-weight model. By removing the need for licensed desktop software, high-performance local hardware, or paid API access, the platform lowers the physical, computational, and financial barriers that can exclude trainees in lower-resourced settings.

All source code, the reference FCSV files, the tutor’s system prompt, and the analysis and figure-generation scripts are released under GPL-3.0. The learning template is pulled from TemplateFlow (Ciric et al., 2022) with server-side caching, so the exact image a learner trains on is a versioned, citable artifact; likewise, the rater-reliability prior that calibrates feedback is regenerated by a released script from the public AFIDs-data. Because the platform is model-agnostic and self-hostable, an institution can reproduce the entire tutoring environment from the public repository.

Beyond the standard anatomical fiducials, the platform’s training architecture is extensible to other landmark-based tasks like deep brain stimulation targets, surgical-resection planning landmarks, or seed points for image segmentation.

This extensibility also makes the validator well suited to multi-site rater qualification: research groups can establish quantitative proficiency benchmarks, and exportable session reports serve as verifiable documentation of a rater’s readiness to contribute to shared research datasets.

### 4.2 Limitations

The guided mode currently uses one template and one contrast, so it does not yet expose learners to pathological anatomy or acquisition variability. Planned directions include additional templates and contrasts, spaced-repetition review of previously missed landmarks, an instructor view for cohort progress and expansion to other landmarks and tasks (e.g., image segmentation).

### 4.3 Evaluation framework and future directions

This platform’s learning-outcome validation is prospective. The guided mode is engineered from established learning-science principles, but we have not yet measured its effect on learner performance. The ORCID-linked opt-in database is designed to accumulate the longitudinal placement data needed for such studies, and we outline the intended evaluation so that it can be replicated:

- Pre/post accuracy on held-out landmarks and templates, comparing guided learning to self-study and to atlas-only instruction;
- Skill transfer from the training template (MNI152NLin2009cAsym T1w) to individual-subject MRI with variable anatomy;
- Reliability convergence (ICC vs. expert consensus) as a function of practice, benchmarked against the trained-rater distribution of Taha et al. (2023);

The platform also fills a specific gap in the open-neuroimaging ecosystem. Tools such as fMRIPrep, MRIQC, and TemplateFlow have standardized how images are processed and shared, but the human skill of anatomical localization that underlies registration and quality control has lacked an equivalent open, teachable resource. Positioned alongside these pipelines, the AFIDs-Validator could be incorporated into graduate curricula, lab onboarding, or multi-site study start-up as a standardized, low-cost proficiency step which complements didactic atlas instruction with hands-on, feedback-driven practice.

## Code and Data Availability

All source code, reference FCSV files, the tutor system prompt, and the analysis and figure-generation scripts (analyze_afids_templates.py, make_figures.py, compute_reliability.py, and the derived rater_reliability.json) are available at https://github.com/afids/afids-validator under GPL-3.0; Figures 4–7 regenerate deterministically from the released templates and placements. The platform is live at https://validator.afids.io. The AFIDs multi-rater dataset (Taha et al., 2023) is at https://github.com/afids/afids-data. The AFIDs protocol is at https://afids.github.io/afids-protocol/.

## Acknowledgements

We thank the TemplateFlow team for maintaining the public template infrastructure, the NiiVue developers (C. Rorden and J. C. Taylor) for the open-source WebGL2 viewer, and the raters who contributed to the AFIDs multi-rater dataset. This research was supported by the Canadian Institutes of Health Research (CIHR) and the Natural Sciences and Engineering Research Council of Canada (NSERC).

## Author Contributions

Conceptualization — J.C.L., A.T.; Software — A.T., J.K., T.K., G.G., J.C.L. (validation engine and reference-template pipeline), A.T. (AI-guided learning mode and rater-reliability calibration), and D.B., A.Th., J.T., O.S., P.P., M.S. (platform development and testing); Formal analysis and Visualization — A.T.; Investigation — A.T., D.B., A.Th., J.T.; Validation — D.B., A.Th., J.T., M.A., A.T.; Data curation — A.T., J.K., T.K., G.G., M.A., V.L., B.M., A.R.K., J.C.L.; Writing – original draft — A.T., J.C.L.; Writing – review & editing — all authors; Supervision — J.C.L., A.R.K.; Funding acquisition — J.C.L., A.R.K.

## Ethics

The neuroimaging data and anatomical fiducial placements underlying the AFIDs-Validator were obtained from openly available brain templates and the AFIDs multi-rater resource (Taha et al., 2023), whose original acquisition was approved by the relevant institutional research ethics boards and, where applicable, conducted with informed participant consent. The platform requires no personal or identifiable information to use; optional ORCID-based sign-in and contribution of placement sessions to the institutional database are strictly opt-in, and no session data are stored without explicit user consent.

